# Seed-dispersal networks in tropical forest fragments: area effects, remnant species, and interaction diversity

**DOI:** 10.1101/701730

**Authors:** Carine Emer, Pedro Jordano, Marco A. Pizo, Milton C. Ribeiro, Fernanda R. da Silva, Mauro Galetti

## Abstract

Seed dispersal interactions involve key ecological processes in tropical forests that help to maintain ecosystem functioning. Yet this functionality may be threatened by increasing habitat loss, defaunation and fragmentation. However, generalist species, and their interactions, can benefit from the habitat change caused by human disturbance while more specialized interactions mostly disappear. Therefore changes in the structure of the local, within fragment, networks can be expected. Here we investigated how the structure of seed-dispersal networks changes along a gradient of increasing habitat fragmentation. We analysed 16 bird seed-dispersal assemblages from forest fragments of a biodiversity-rich ecosystem. We found significant species-, interaction- and network-area relationships, yet the later was determined by the number of species remaining in each community. The number of frugivorous bird and plant species, their interactions, and the number of links per species decreases as area is lost in the fragmented landscape. In contrast, network nestedness has a negative relationship with fragment area, suggesting an increasing generalization of the network structure in the gradient of fragmentation. Network specialization was not significantly affected by area, indicating that some network properties may be invariant to disturbance. Still, the local extinction of partner species, paralleled by a loss of interactions and specialist-specialist bird-plant seed dispersal associations suggests the functional homogenization of the system as area is lost. Our study provides empirical evidence for network-area relationships driven by the presence/absence of remnant species and the interactions they perform.

**RESUMO:** Interações de dispersão de sementes formam um processo ecológico chave em florestas tropicais onde colaboram na manutenção do funcionamento do ecossistema. Porém, esta funcionalidade pode estar ameaçada pelo aumento na perda e fragmentação do habitat. Enquanto espécies generalistas e suas interações podem se beneficiar da mudança de habitat causada por distúrbios antrópicos, interações envolvendo espécies mais especialistas são, na maioria, eliminadas. Desta forma, mudanças nas redes locais, dentro de fragmentos florestais, são esperadas. Neste trabalho nós investigamos como a estrutura de redes de dispersão de sementes mudam em um gradiente de fragmentação do habitat. Nós analisamos 16 comunidades de dispersão de sementes espacialmente explícitas e distribuídas em fragmentos florestais de um ecossistema rico em biodiversidade. Nós encontramos relações significativas entre a área do fragmento e espécies, interações e estrutura das redes, sendo que o último foi determinado pelo número de espécies remanescentes em cada comunidade. O número de espécies de aves frugívoras e plantas e as interações entre eles, bem como o número de links por espécie diminuíram significativamente conforme a área dos fragmentos é perdida. Por outro lado, o aninhamento da rede mostrou uma relação negativa com a área do fragmento, sugerindo um aumento da generalização da estrutura das redes com a fragmentação do habitat. No entanto, o grau de especialização das redes não foi afetado pela área, indicando que algumas propriedades de rede podem ser resistentes à perturbação. Sendo assim, a extinção local de espécies parceiras, conjuntamente com a perda das interações e associações planta-dispersor mais especializadas, sugere uma homogeneização do sistema conforme a área do fragmento é perdida. Nosso estudo fornece evidências empíricas para as relações rede-área, sendo estas direcionadas pela presença e/ou ausência das espécies remanescentes bem como das interações que estas realizam.

## INTRODUCTION

### THE ON-GOING, FAST-PACED DEFORESTATION HAS CREATED FRAGMENTED FORESTS POSING INCREASING

challenges to the conservation of species interactions and ecological processes (Janzen 1974, Bruna et al. 2005, McConkey et al. 2016, Tylianakis & Morris 2017). Even though fragmentation is well documented to affect species persistence and the connectivity among forest patches (da Silva & Tabarelli 2000, Rybicki & Hanski 2013, Hagen et al. 2012), the remnant fragments are also proven to hold an important fraction of biodiversity (Barlow et al. 2007, Sekercioğlu et al. 2007, Bongers et al. 2015). The pool of species and interactions persisting in forest fragments is formed by a non-random subset of smaller-sized species while the whole spectrum of sizes found at the landscape level can only occur in large, more pristine areas (Morante-Filho et al. 2015, Emer et al. 2018). Such local assemblages configure local networks of interacting partner species whose structure may be affected by the constraints imposed by fragment area (Aizen et al. 2012, Emer et al. 2013, Bomfim et al. 2018) while resilience to habitat loss may occur in specific systems (e.g., Passmore et al. 2012). A recent theoretical framework has proposed different mechanisms by which network-area relationships may emerge, such as the constraints imposed by species-area relationships, the selection towards generalist species and the dispersal limitation in smaller spatial scales (Galiana et al. 2018). Yet, whether network structure is robust enough to be maintained across fragmented forests, or whether it changes as habitat area is lost is still an open question to be tested with empirical data.

Ecological networks are the outcomes from associations among interacting species (Jordano et al. 2003, Lewinshon & Prado 2006). Mutualistic associations between animal and plants are a fundamental process to maintain forest dynamics (Herrera & Pellmyr 2002, Dennis et al. 2007). Seed dispersal by birds, in which both interacting species benefit by using resource from one another, are ubiquitous in tropical forests in which up to 90 percent of the plant species rely on birds to be effectively dispersed (Howe & Smallwood 1982, Jordano 1995). Such associations between tropical frugivorous birds and the plants they feed upon can form complex and diverse seed-dispersal networks, characterized by a large number of species and interactions (Bascompte & Jordano 2007, Schleuning et al. 2014, Dügger et al. 2018). Disruptions in the functionality of this system can be inferred by the emergence of ecological patterns depicted from the correct interpretation of the structural properties of the networks (Bascompte & Jordano, 2007, Memmott 2009, Howe 2016). For instance, changes in the mean number of interactions (links) per species may indicate a reduction on the number of partners available to interact with in smaller areas, in which generalist species are favoured (Morante-Filho et al. 2016, Galiana et al. 2018). Likewise, increases in nestedness, a pervasive pattern in mutualistic networks that suggests the presence of highly generalist species interacting with both generalist and specialist ones, may suggest the loss of specialist interactions in disturbed communities (Aizen et al. 2012, Bomfim et al. 2018). Similarly, a reduction of network specialization, estimated by the H2’ index based on Shannon diversity, would suggest the community is losing diversity of interactions among species, with the prevalence of more generalist species.

Thus, the winner species and their winner interactions able to thrive despite the disturbances imposed by habitat loss and fragmentation may be the ones defining the configuration of the local networks (Hobbs et al. 2006, Tabarelli et al. 2012, Vidal et al. 2014). For instance, diet generalist species are expected to be more resilient to habitat change due to their ability to feed on, and to be dispersed by, a wider range of interacting partners (Tabarelli et al. 2012, Morante-Filho et al. 2015). Likewise, generalist small-bodied bird species and plant species typical of secondary forest may form loose, generalists associations in fragmented landscapes (Emer et al. 2018), which may result in an increased generalization in seed dispersal networks. Furthermore, habitat fragmentation generates changes on biotic and abiotic conditions (Murcia 1995, Laurance et al. 2007) such as increases in temperature and light conditions that may favour the competitive ability of some plant species over the others (Laurance et al. 2010, Sfair et al. 2016). Associated to the negative effects of habitat reduction lays defaunation, a pervasive phenomenon characterized by the local or functional extinction of a given species, generally towards the larger body mass ones and often related to the reduction of habitat area (Dirzo et al., 2014, Young et al. 2016). Thus, interactions performed by large-bodied species dispersing large-seeded plants are expected to be absent in smaller areas (Pérez-Mendez et al. 2016, Emer et al. 2018) therefore affecting the local, within-fragment network structure.

Here we aim to understand how the reduction of habitat through landscape fragmentation affects the structure of tropical seed-dispersal networks. We gathered data of avian-seed dispersal interactions located in 16 fragments of the Atlantic Forest, a highly diverse and threaten tropical biome (Ribeiro et al. 2009). Our first prediction is that the number of species present in each fragment will follow a species-area relationship (MacArthur & Wilson 1967, Rybicki & Hanski 2013), i.e., the number of species reduces according to the loss of habitat area. Secondly, because smaller area fragments are disturbed habitats inhabited mostly by generalist species (Morante-Filho et al. 2016, Emer et al. 2018) able to cope with the biotic and abiotic changes promoted by edge-effects associated to other human-disturbance (Laurance et al. 2007), we expect a positive relationship between fragment area, the total number of bird-seed dispersal interactions and the mean number of links per species. Because the number of bird-seed dispersal interactions in forest fragments is selectively reduced towards generalist species (Morante-Filho et al. 2016, Emer et al. 2018), we expect that the remaining generalist interactions will lead to changes in network structure towards higher nestedness and lower specialization, with interaction links associated to a few dominant species.

## METHODS

### DATASET

We compiled 16 studies of avian plant-frugivore interactions sampled at the community level along the Atlantic Forest biome, a hotspot of biodiversity highly threaten by increasing human pressure (Ribeiro et al. 2009, Joly et al. 2014; Fig. 1, Table S1). The study areas varied from 0.66 to 42,000 ha, along a gradient of disturbance from semi-pristine Biological Reserves and State Parks to secondary forest fragments and restored private lands. The matrix surrounding the fragments is variable, including sugar cane fields, crop plantations, secondary forest, and urban areas. Overall the studies included here originally aimed to record bird-eating-fruit interactions and not necessarily effective seed dispersal; therefore we carefully checked every dataset and removed any interaction that would not characterize plant dispersal events, such as seed predation. In the particular case of parrots we excluded them from the analysis despite some rare evidences pointing to their role in effective seed dispersal of large nuts (Tella *et al.* 2016); in most cases we could not establish in the original papers whether the frequency of interactions involving parrots actually implied legitimate dispersal. We updated and standardized plant and bird species names using the taxize package (Chamberlain & Szocs, 2013) in R (R Development Core Team 2014).

**FIGURE 1.**
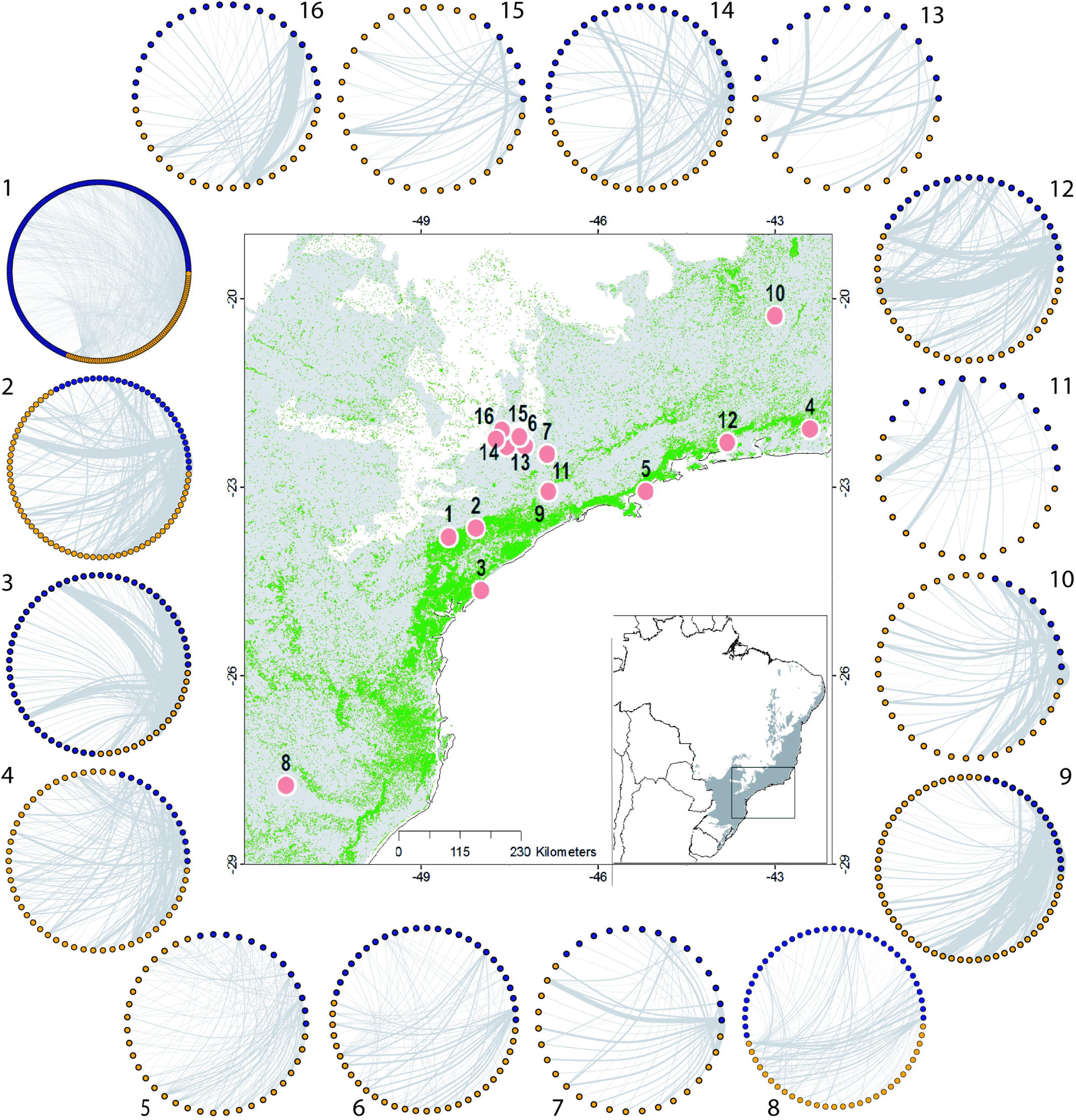
The structure of the 16 avian seed-dispersal networks and their location in the Atlantic Forest, SE Brazil. In each network, blue dots represent plant species, orange dots bird species, and the grey lines are the seed-dispersal interactions among them; the width of the grey lines correspond to the frequency of interactions in each area. Communities are numbered according to a decreasing order of fragment area (see Table S1 for details of each site): 1- Parque Estadual (PE) Intervales; 2- PE Carlos Botelho; 3- PE Ilha do Cardoso; 4- PE Poço das Antas; 5- PE Ilha Anchieta; 6-Minas Gerais fragment; 7- Estação Ecológica de Aracuri; 8- Mata Santa Genebra; 9- Itatiba fragment; 10- Rio Claro fragment; 11- restored area 15 years old; 12- restored area 25 years old; 13- restored area 57 years old; 14- São Paulo fragment; 15- restored area 8 years old; 16- Rio de Janeiro fragment.

### NETWORK METRICS

For each community we built a correspondent seed-dispersal network as a bipartite graph represented by a weighted matrix *A*_*ij*_ in which the rows represent the *i* plant species and columns represent the *j* bird species. The elements *a*_*ij*_ indicate the frequency of interactions between the plant species *i* and frugivore bird species *j*. Changes in network structure were assessed by estimating the following non-correlated metrics (r < 0.7):

i. the number of bird species (*A*);
ii. the number of plant species (*P*);
iii. the number of interactions (*I*);
iv. the mean number of links per species (*L*);
v. weighted nestedness (*wNODF*), which estimates the presence of generalist species interacting with more specialist ones, based on the NODF index (Almeida-Neto et al. 2008);
vi. H2’ specialization (*H2’*), which estimates how strong species partition their interaction partners. It measures the deviation of non-specialization in which values close to zero means low specialization of the interacting species in the community while values close to one indicate high levels of specialization (Blüthgen et al. 2006).

### LANDSCAPE METRICS

Landscape configuration can play an important role in determining species persistence in fragmented landscapes (Ribeiro et al. 2009) and as such may influence NAR’s as well. Therefore, for each studied community we calculated the fragment area (ha), the functional connectivity, and the average isolation. We used the same remnant forests map generated by Ribeiro et al. (2009) as input map for the calculation of the landscape metrics. The maps had 50 m spatial resolution for Albers Equal Area projection and South America Datum (SAD 69). Because some birds can successfully move among fragments within a given distance (Pizo et al. 2011; Cornellius et al. 2017; Vélez et al. 2015), and some interactions functionally persist throughout a fragmented landscape (Emer et al. 2018), we estimated the functional connectivity as the amount of forest (ha) that any species can have access if it is able to cross a 500 m matrix (Ribeiro et al., 2009). Average isolation was calculated by randomly choosing 1000 points in a 5 km buffer around the sampling coordinates and extracting the Euclidean mean distance (in meters) to the nearest forest fragment.

### DEFAUNATION

Likewise, for each studied community we estimated the intensity of defaunation relatively to the regional pool of species expected for the Atlantic Forest biome. To do so, we calculated the difference between the sum of all bird body masses at the regional landscape scale (considering all frugivore bird species present in the Atlantic-Frugivory database [Bello et al 2017] as a proxy for the regional pool of species) and the summed body masses of the birds that were recorded interacting with fruits in each fragment. Defaunation is based only on the presence/absence of species in a given fragment.

### STATISTICAL ANALYSES

To understand whether and how the structure of avian seed-dispersal networks is affected by habitat change we used Linear Regressions and General Least Square (gls) models in the nlme package (Pinheiro et al., 2016). Individual models were fit for each network metric therefore considering the number of plant species, the number of bird species, the number of interactions, the mean number of interactions per species, nestedness and H2’ specialization as response variables. Network metrics were logit-transformed to solve the issue of being bounded from zero to one, when needed (Warton & Hui, 2011). Because defaunation is negatively correlated with fragment area (r = - 0.89) and the three landscape metrics are highly correlated among them (Table S2), we parsimoniously chose area as the variable representing changes in habitat configuration. Therefore, fragment area was fitted as the independent variable along with two covariates: (i) sampling intensity, calculated as the root square of the number of interactions divided by the root square of the network size (sum of plant species and bird species), to control for differences in sampling effort among studies (following Schleuning et al. 2012), and (ii) forest type, classified according to the vegetation structure of each community (Ombrofilous Forest, Semidecidous Forest, Araucaria Forest, and restored forest) to control for possible effects of biotic and abiotic conditions intrinsic to each forest formation. Our spatially explicit models and model selection follows the protocols suggested in Zuur et al. (2009). We did not detect spatial correlation structure in our models after accounting for the geographic coordinates of each study site in the gls models, according to the Akaike Information Criterion (AICc). Therefore, we fit the full model to linear regressions with ‘Maximum Likelihood estimation’ while applying the ‘dredge’ function (Barton 2013) to select for the best-fixed structure according to AICc. ‘Sampling intensity’ was used as a fixed parameter to test for the effects of area and forest type solely. Then, we used the Akaike weight of evidence (wAICc) to obtain the relative importance of the different models (Burnham and Anderson 2002). Finally, we present the coefficients of each predictor and the variance explained by the optimal linear model. Lastly, we used two different but complementary null models to test for changes in the number of links per species, nestedness and H2’ specialization independently of network size by fitting standardized z-scores in the same model structure described above. Both null models constrained network size and were created using the ‘nullmodel’ function. First, we used ‘method = 1’ (Patefield algorithm) to randomize the distribution of links among species while maintaining the marginal totals constant, i.e., species have the same total number of interactions in the observed and randomized matrices, therefore maintaining their biological characteristic of being common or rare in the community (Dorman et al. 2009). Second, we used ‘method = 3’ (vaznull algorithm) to randomize the number of interactions among species without constraining marginal totals but maintaining connectance constant, i.e., species have the same number of qualitative interactions as in the observed matrix (Vázquez et al. 2007). Then, the mean and standard deviation of 100 iterations of each null model were contrasted against the observed values of links per species, nestedness and specialization; the resultant z-scores were used as the response variable in the statistical models. Network metrics and null models were estimated in the bipartite package (Dormann et al., 2009). All analyses were run in R (R Development Core Team, 2014).

## RESULTS

Our dataset comprises 281 plant species (plus 54 plants identified and/or morphotyped at the genus or family level) that interact with 175 frugivore bird species. We recorded a total of 7637 interactions and 2389 unique pairwise links, in which a bird species dispersed the seeds of a plant species in the studied fragments of the Atlantic Forest.

The mean frequency of interactions per fragment (472.4 ± 363.4) was 16 times greater than the mean number of bird (32.3 ± 17.9) and plant species (30.7 ± 42.6) recorded in each study site. The most abundant interactions were those performed by generalist bird species dispersing secondary-forest plant species, such as *Thraupis sayaca* dispersing seeds of *Cecropia pachystachya* (61 events of seed-dispersal between three fragments; Fig S1). However, most of the events of seed-dispersal by frugivore birds were rare given that 64% of the interactions occurred only once across the studied fragments (Fig. S1).

We found significant species, interaction- and network-area relationships (Fig. 2, Table 1). First, the number of bird and plant species decreased significantly with fragment area. Second, both the number of bird seed-dispersal interactions and the mean number of links per species were negatively affected by the reduction of fragment area. And finally, network nestedness showed a positive and significant relationship with increasing fragmentation when controlling for sampling intensity while specialization showed a tendency to decrease as area is lost, even though the variance explained by this model was quite low. When network size was controlled for by the use of the null models, only sampling intensity explained changes in the mean number of links per species, network nestedness and H2’ specialization (Table S3).

**TABLE 1.**
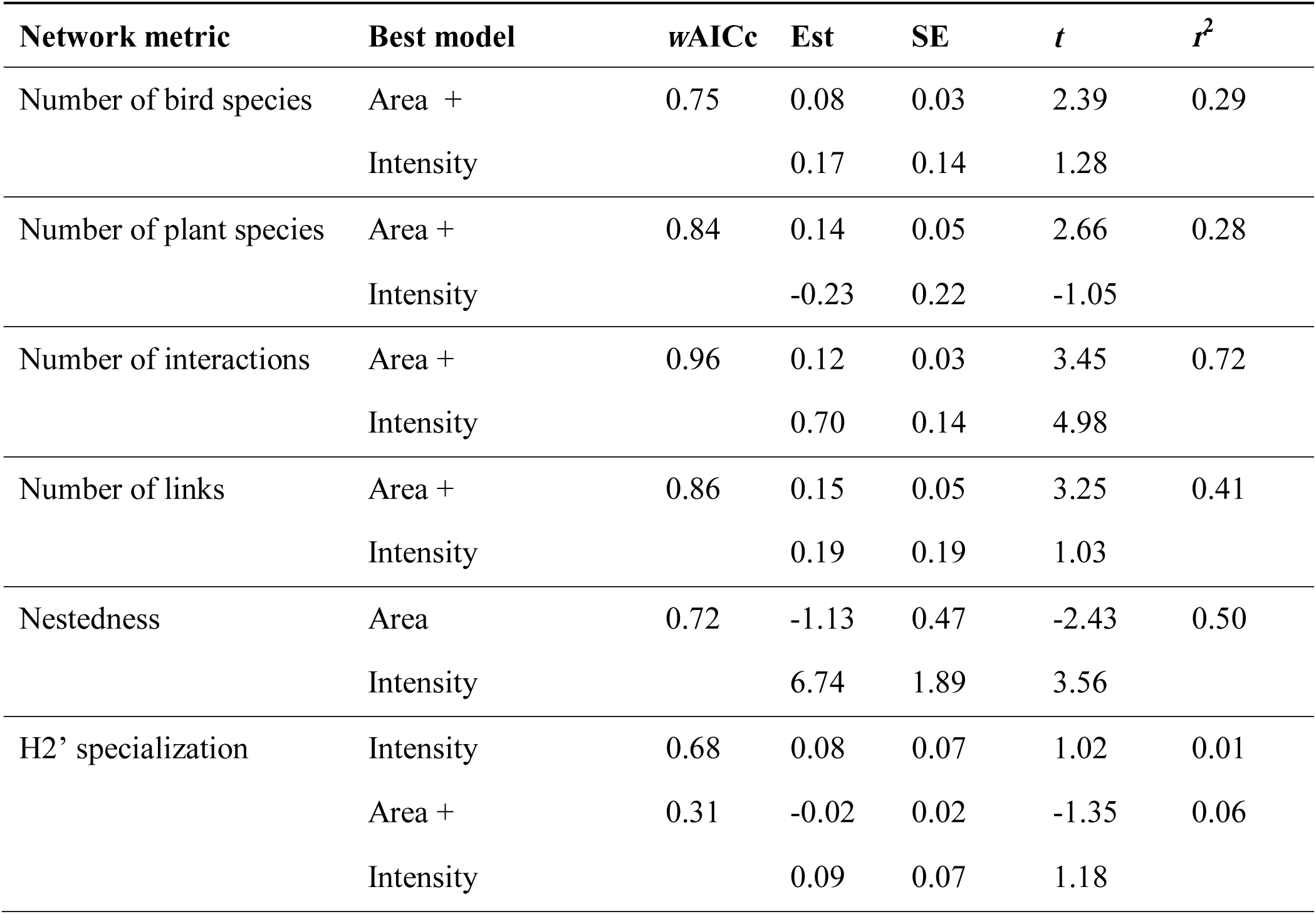
Results of the Linear Regression Models after AICc model selection testing the effects of fragment area on the structure of avian seed-dispersal networks. Area corresponds to the logarithmic scale of hectares per fragment. Intensity corresponds to the average number of interactions per species and was used as a fixed parameter to control for differences in sampling effort among studies. Only models with delta AICc < 2 were selected as plausible explanations for the observed patterns (see Table S1 for full model selection). *w*AICc gives an estimate of the probability of that given model to be the best choice under the AICc criteria. *r*^2^ gives an estimation of the variance explained by the optimal model while the coefficient *t* corresponds to the importance of that parameter within the model (*t* > 2 indicates the coefficient is significant with > 95% confidence). ‘+’ indicates that the next presented variable is co-varying to explain the changes on the corresponding network metric in the corresponding model.

**FIGURE 2.**
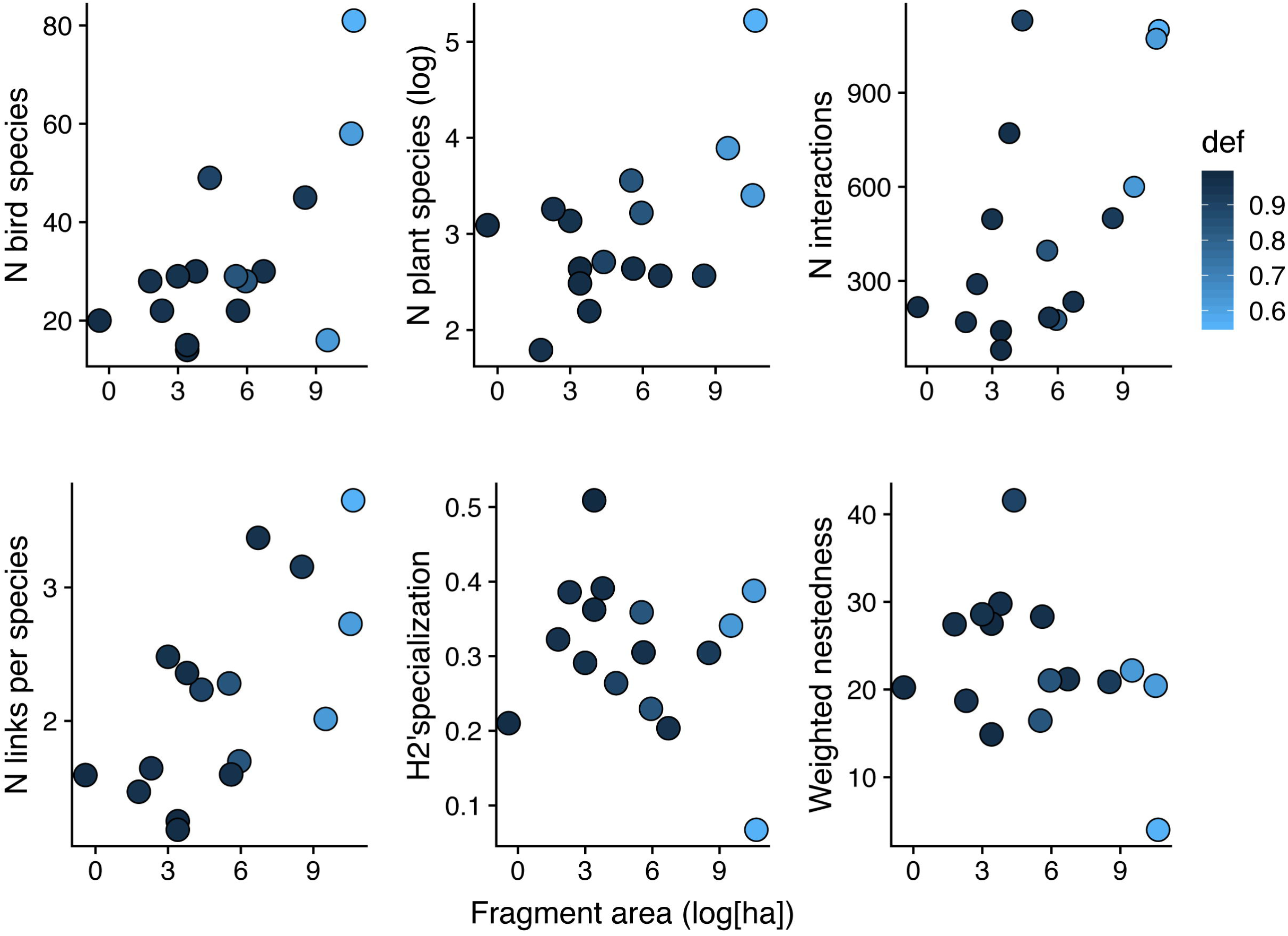
The effects of fragment area on the structure of seed-dispersal networks of the Atlantic Forest. Each point corresponds to a single forest fragment. The gradient of colours correspond to the intensity of defaunation (‘def’) in each fragment, calculated as the difference between the total body mass of the frugivore birds expected for the Atlantic Forest (based on Bello et al. 2017) and the total body mass of bird frugivores recorded in each network. Defaunation and fragment area are highly correlated (r = - 0.89).

## DISCUSSION

We found significant changes of network structure following the lost of habitat area in the fragments of the Atlantic Forest. The direction of these changes is driven by the non-random loss of frugivorous bird and plant species and the interactions they perform. Such changes derived from the lower number of links per species and the higher network nestedness observed in smaller area fragments, in which bird-seed dispersal interactions performed by small-sized generalist species are predominant (Emer et al. 2018). Yet, the reduced community size in smaller areas led to the structural changes observed on the organization of seed-dispersal interactions, depicted by the influence of sampling intensity and the lack of significant effects when tested against proper null models. Nonetheless, network specialisation seems to be an invariant property of those communities given that it was not affected by habitat change or pairwise species losses.

Bird seed-dispersal networks of the Atlantic Forest are losing species and interactions as the landscape becomes more fragmented. Beyond the potential consequences for network stability (May 1972, Valdovinos 2019) detected by the increasing nestedness and reduced number of links performed by the remnant species, those local communities are likely to be also losing important ecological functions. Species- and interaction-richer communities mostly hold higher functional diversity while the impoverished ones are likely to have gaps on ecosystem functionality (Isbell et al., 2011; Saavedra et al. 2014, Montoya et al., 2015). The impoverishment of plant species in forest remnants from the Northeast of the Atlantic Forest has led to a biotic homogenization across the fragmented landscape (Lobo et al. 2011). Likewise, the impoverishment of interactions found in this study and the persisting interactions performed by smaller-sized generalist species in the fragmented landscape (Emer et al. 2018) suggests that the Atlantic Forest is losing specific ecological functions, which may trigger medium to long-term functional homogenization of interactions throughout the landscape (Olden et al. 2004, Laliberté & Tylianakis 2010). The changes on the number of links per species according to habitat fragmentation may support our *a posteriori* hypothesis of functional homogenization potentially driven by the persistence of a few, functionally-redundant sub-groups of interacting species that are able to cope with the unstable conditions found in most forest fragments (Murcia 1995, Harper et al. 2005, Passmore et al. 2012).

The increasing nestedness of the bird-seed dispersal networks indicates that communities in smaller area fragments predominantly include a core of generalist species that interact with a sub-group of more specialist ones, while there is a lack, or at least a reduction of, specialist-specialist interactions. In other words, the reduced species pool on smaller fragments are likely to promote their generalization in terms of mutualistic partners while using whichever resource is available in these more disturbed areas. Thus, it may indicate a community-level shift towards more generalist interactions as fragmentation increases (Hobbs et al. 2006; Kiers et al 2010). Likewise, changes in the behaviour of the species that persist in smaller fragments, or fluctuations in population dynamics, could also explain the increasing network generalization (Awade et al. 2017; Cornelius et al. 2017). Furthermore, the reduced community size in smaller areas apparently leads to some structural changes in the organization of seed-dispersal interactions, as suggested by the influence of sampling intensity on network nestedness when tested against proper null models.

In turn, network specialization was not significantly affected by habitat reduction. This suggests that, despite negative effects of fragmentation on network size, the way remnant species interact in terms of partners diversity is similar between small assemblages of bird and plant species in smaller areas, and large assemblages in larger areas. Considering that the minimum value of the H2’ index indicates maximum niche overlap among interacting species (while the opposite indicates maximum niche divergence; Schleuning et al. 2012), our analysis reveals that niche partitioning of fruit food resources provided by the plants to the birds, or the seed dispersal service promoted by the birds to the plants, are not affected by the loss of area in smaller sized fragments. This suggests that network specialization in terms of niche partitioning is driven by different mechanisms other than habitat area (Schleuning et al. 20144, Mello et al. 2015). The presence of naturalized alien species in the system, mainly plants such as *Melia azedarach* and *Psidium guajava*, may numerically compensate the absence of some native ones, which are more sensitive to habitat change. Yet, their functionality as food resources for the bird community appears impoverished.

Tropical forest fragments are hyperdynamic systems facing population and community changes in response to primary effects of habitat loss followed by subsequent edge effects and stochastic events (Laurance et al. 2002, Laurance et al. 2010). Furthermore, the Atlantic Forest landscape holds very high levels of beta-diversity and turnover of birds, plants, and their interactions (Morante-Filho et al. 2016, Farah et al. 2017, Emer et al. 2018), which could lead, at least partially, to the observed network-area relationships. Such community heterogeneities can be further influenced by other anthropogenic impacts such as hunting and logging, whose frequency and intensity of occurrence are context-dependent and vary among fragments (Joly et al. 2014, Galetti et al. 2016). Therefore, long-term effects of habitat fragmentation on network structure may go through different trajectories (Thompson 2005) but the impoverishment of the system is unlikely to change.

Finally, our findings have important implications for advancing the understanding of network-area relationships (NARs, *sensu* Galiana et al. 2018). While species-area relationships are one of the most pervasive rules in ecology (MacArthur & Wilson 1967, Lawton 1999), empirical data demonstrating whether or not interactions- and network-area follow the same rules are rare (but see Aizen et al. 2012, Passmore et al. 2012, Emer et al. 2013, Bomfim et al. 2018 for examples on mutualistic networks). Our results showed that, as expected, bird and plant species as well as their interactions, were strongly affected by fragment area. Likewise, the number of links per species and nestedness also changed with area but not network specialization. Yet, when the number of species in each community was considered using the null models, none of the network properties changed with area, only with sampling intensity. These findings suggest that area *per se* does affect network structure, but primarily mediates the presence/absence of remnant species in the fragments, as well as the interactions they perform. Furthermore, the lack of significant changes in network specialization independently of local species richness, contrasting with the richness-associated changes of the previous metrics, may suggest that the stability of complex system depends on the specific parameter and the system analysed (May 1972, Thébault & Fontaine 2010, Valdovinos et al. 2019). Specialization of the seed-dispersal interactions could be seen as an invariant property of the networks, at some extent robust to habitat loss. On the other hand, network size and the consequent number of interactions and nestedness seem to be more prone to collapse as disturbance increases with landscape fragmentation and reduced habitat area. Therefore, the loss of species and their interactions following habitat loss may be a good predictor of the directions of changes in network structure, providing guidance for an overall theory of NARs.

## ACKNOWLEDGMENTS

We thank Carolina Bello to provide the map used in Figure 1 and Carolina Carvalho for statistical advice. This work was supported by the Fundação de Amparo à Pesquisa do Estado de São Paulo (BIOTA – FAPESP 2014/01986-0). CE was supported by a postdoctoral fellowship (FAPESP 2015/15172-7). MG and MAP received a fellowship from the National Council for Scientific and Technological Development (CNPq). MG and PJ were funded in part by CYTED (project 418RT0555).

## DATA AVAILABILITY STATEMENT

Upon acceptance, the dataset used in this manuscript will be made available on the main author’s GitHub repository (https://github.com/carineemer/network_area).

## SUPPLEMENTARY INFORMATION

Content

Table S1-S4.

Fig. S1

