## Supplementary material for "Seed-dispersal networks in tropical forest fragments: area effects, remnant species, and interaction diversity": SM

Biotropica

**Content**

Table S1 to S4

Fig S1

**Table S1.** Description of the bird-seed dispersal interaction data used in this study. All studies were carried out in the Atlantic Forest biome, SE Brazil. Forest types correspond to: OF = Ombrofilous Forest, SDF = Semideciduous Forest, AF = Araucaria Forest, RF = Restored Forest. Ni = total number of interactions; SI = sampling intensity; L = mean number of links per species; C = connectance; *w*NODF = weighted nestedness; Ev = interaction evenness; Mod = weighted modularity; Def = defaunation index; SF = sampling focus, which corresponds to: P = plant oriented, interactions sampled based on focal observations; A = animal oriented, interactions sampled based on diet analyses (e.g., faeces sampling). See methods in the main text for details of each network parameter. Only non-correlated network metrics were used in the statistical analyses, r < 0.7)

| **ID*** | **Lat** | **Long** | **Forest Type** | **Area (ha)** | **N bird species** | **N plants species** | **Size** | **Ni** | **SI** | **L** | **C** | ***w*NODF** | **H2** | **Ev** | **Mod** | **Def** | **SF** |
| --- | --- | --- | --- | --- | --- | --- | --- | --- | --- | --- | --- | --- | --- | --- | --- | --- | --- |
| 1 | -24.316965 | -48.387175 | OF | 42000 | 81 | 185 | 266 | 1100 | 2.03 | 3.65 | 0.06 | 4.00 | 0.07 | 0.71 | -0.70 | 0.557 | PA |
| 2 | -24.131389 | -47.949167 | OF | 37644 | 58 | 30 | 88 | 1072 | 3.49 | 2.73 | 0.14 | 20.43 | 0.39 | 0.66 | -0.58 | 0.615 | PA |
| 3 | -25.127786 | -47.957463 | OF | 13500 | 16 | 49 | 65 | 600 | 3.04 | 2.02 | 0.17 | 22.17 | 0.34 | 0.62 | -0.59 | 0.626 | P |
| 4 | -22.53629 | -42.277658 | OF | 5052 | 45 | 13 | 58 | 500 | 2.94 | 3.16 | 0.31 | 20.87 | 0.30 | 0.77 | -0.66 | 0.926 | P |
| 5 | -23.548369 | -45.062541 | OF | 828 | 30 | 13 | 43 | 234 | 2.33 | 3.37 | 0.37 | 21.18 | 0.20 | 0.85 | -0.76 | 0.958 | P |
| 6 | -20.802611 | -42.858746 | SDF | 380 | 28 | 25 | 53 | 176 | 1.82 | 1.69 | 0.13 | 21.04 | 0.22 | 0.65 | -0.62 | 0.838 | P |
| 7 | -28.223141 | -51.166857 | AF | 272 | 22 | 14 | 36 | 184 | 2.26 | 1.60 | 0.19 | 28.31 | 0.31 | 0.64 | -0.61 | 0.96 | P |
| 8 | -22.818101 | -47.113757 | SDF | 250 | 29 | 35 | 64 | 397 | 2.49 | 2.28 | 0.14 | 16.46 | 0.36 | 0.67 | -0.58 | 0.837 | P |
| 9 | -22.943263 | -46.749949 | SDF | 80 | 49 | 15 | 64 | 1130 | 4.20 | 2.23 | 0.19 | 41.59 | 0.26 | 0.63 | -0.70 | 0.897 | PA |
| 10 | -22.480979 | -47.592293 | SDF | 44 | 30 | 9 | 39 | 771 | 4.45 | 2.36 | 0.34 | 29.78 | 0.39 | 0.67 | -0.56 | 0.963 | P |
| 11 | -22.825234 | -47.427761 | RF | 30 | 14 | 14 | 28 | 141 | 2.24 | 1.25 | 0.18 | 27.52 | 0.51 | 0.58 | -0.49 | 0.986 | PA |
| 12 | -22.671644 | -47.204638 | RF | 30 | 15 | 12 | 27 | 80 | 1.72 | 1.19 | 0.18 | 14.88 | 0.36 | 0.55 | -0.63 | 0.971 | PA |
| 13 | -22.568342 | -47.504987 | RF | 20 | 29 | 23 | 52 | 497 | 3.09 | 2.48 | 0.19 | 28.58 | 0.29 | 0.65 | -0.70 | 0.95 | PA |
| 14 | -23.545856 | -46.721177 | SDF | 10 | 22 | 26 | 48 | 290 | 2.46 | 1.65 | 0.14 | 18.72 | 0.39 | 0.61 | -0.56 | 0.952 | P |
| 15 | -22.708708 | -47.610207 | RF | 6 | 28 | 6 | 34 | 169 | 2.23 | 1.47 | 0.30 | 27.44 | 0.32 | 0.69 | -0.62 | 0.956 | P |
| 15 | -22.767379 | -43.694394 | SDF | 0.66 | 20 | 22 | 42 | 217 | 2.27 | 1.59 | 0.15 | 20.23 | 0.21 | 0.58 | -0.81 | 0.971 | P |

*Identification of the networks used in the study with corresponding references.

1. Silva, W. R., De Marco, P., Hasui, E., and Gomes, V.S.M. (2002). Patterns of fruit-frugivores interactions in two Atlantic Forest bird communities of South-eastern Brazil: implications for conservation. In: Seed dispersal and frugivory: ecology, evolution and conservation (eds. Levey, D.J, Silva, W.R, and Galetti, M.). Wallinford: CAB International. pp. 423-435.
2. Rodrigues, S.B.M. (2015). Rede de interações entre aves frugívoras e plantas em uma área de Mata Atlântica no sudeste do Brasil. Master thesis. UFSCAR Sorocaba.
3. Castro, E. R. (2015). Fenologia reprodutiva do palmito Euterpe edulis (Arecaceae) e sua influência na abundância de aves frugívoras na Floresta Atlântica. PhD Thesis. UNESP Rio Claro.
4. Correia, J. M. S. (1997). Utilização de espécies frutíferas da Mata Atlântica na alimentação da avifauna da Reserva Biológica de Poços das Antas, RJ. MSc Dissertation.  UNB Brasília.
5. Alves, K. J. F. (2008). Composição da avifauna e frugivoria por aves em um mosaico sucessional na Mata Atlântica. MSc Dissertation. UNESP Rio Claro.
6. Fadini, R.F. and Marco Jr., P. de. (2004). Interações entre aves frugívoras e plantas em um fragmento de Mata Atlântica de Minas Gerais. Ararajuba 12: 97-103.
7. Kindel, A. (1996). Interações entre plantas ornitocóricas e aves frugívoras na Estação Ecológica de Aracuri, Muitos Capões, RS. MSc Dissertation. UFRGS Porto Alegre.
8. Galetti, M., Pizo, M.A. (1996). Fruit eating birds in a forest fragment in southeastern Brazil. Ararajuba 4: 71-79.
9. Pizo, M. A. (2004). Frugivory and habitat use by fruit-eating birds in a fragmented landscape of southeast Brazil. Ornitol Neotrop, 15 (Suppl.): 117–126.
10. Athiê, S. (2009). Composição da avifauna e frugivoria por aves em um mosaico de vegetação secundária em Rio Claro, região centro-leste do estado de São Paulo. MSc Dissertation. UFSCar São Carlos.
11. da Silva, F. R., Montoya, D., Furtado, R., Memmott, J., Pizo, M.A., and Rodrigues, R.R. (2015). The restoration of tropical seed dispersal networks. Rest. Ecol., doi: 10.1111/rec.12242
12. da Silva, F. R., Montoya, D., Furtado, R., Memmott, J., Pizo, M.A., and Rodrigues, R.R. (2015). The restoration of tropical seed dispersal networks. Rest. Ecol., doi: 10.1111/rec.12243
13. da Silva, F. R., Montoya, D., Furtado, R., Memmott, J., Pizo, M.A., and Rodrigues, R.R. (2015). The restoration of tropical seed dispersal networks. Rest. Ecol., doi: 10.1111/rec.12244
14. Hasui, E.  (1994). O papel das aves frugívoras na dispersão de sementes em um fragmento de floresta semidecídua secundária em São Paulo, SP. MSc Dissertation. USP São Paulo.
15. Robinson, V. (2015). Índice de importância das aves como dispersoras de sementes para uma comunidade vegetal reflorestada em Piracicaba. BSc Dissertation, UNESP, Rio Claro.
16. Silva, R.F.de M. (2011). Interações entre plantas e aves frugívoras no campus da Universidade Federal do Rio de Janeiro. Undergrad Dissertation. UFRJ Rio de Janeiro.

**Table S2**. Pearson correlations between the landscape metrics calculated for each study site. Description of each metric is found in the main text.

|  | Area (ha) | Average isolation | Functional connectivity |
| --- | --- | --- | --- |
| Area (ha) | 1 | -0.782 | 0.755 |
| Average isolation | -0.782 | 1 | -0.897 |
| Functional connectivity | 0.755 | -0.897 | 1 |

**Table S3.** Results of the Linear Regression Models after AICc model selection testing for the effects of fragment area on the structure of avian seed-dispersal networks. Standardized z-scores were used to control for potential influence of network size in the observed values. Z-score_1 corresponds to the observed value compared against null models constructed based on the Patefield algorithm while z-score_3 was based on vaznull algorithm (see Methods for details in the main text). Area corresponded to the logarithmic scale of the hectares per fragment. Intensity corresponds to the average number of interactions per species and was used to control for differences in sampling effort among fragments. Only models with delta AICc < 2 were selected as plausible explanations for the observed patterns. *w*AICc gives an estimate of the probability of a given model to be the best choice under the AICc criteria. *r*^2^ gives an estimation of the variance explained by the optimal model while the coefficient *t* corresponds to the importance of that parameter within the model (*t* > 2 indicates the coefficient is significant with > 95% confidence). For the number of links, only z-score with Patefield algorithm was calculated because the vaznull algorithm does not allow changes on the mean number of links per community due to the constrained connectance applied.

| **Network metric** | **Best model** | **AICc** |  | **ΔAICc** | ***w*AICc** | **Est** |  | **SE** | **t** | ***r*^2^** |
| --- | --- | --- | --- | --- | --- | --- | --- | --- | --- | --- |
| Number of links  (z_score_1) | Intensity | 102.4 |  | 0 | 0.86 | -6.87 |  | 1.60 | -4.28 | 0.53 |
| Nestedness  (z-score_1) | Intensity | 69.6 |  | 0 | 0.86 | -2.46 |  | 0.58 | -4.27 | 0.53 |
| Nestedness  (z-score_3) | Intensity | 70.6 |  | 0 | 0.85 | -1.48 |  | 0.59 | -2.50 | 0.26 |
| H2’ specialization (z-score_1) | Intensity | 114.8 |  | 0 | 0.84 | 18.25 |  | 2.37 | 7.72 | 0.80 |
| H2’ specialization (z-score_3) | Intensity | 87.6 |  | 0 | 0.83 | 6.56 |  | 1.01 | 6.48 | 0.73 |

**Table S4.** Results of the Linear Regression Models after AICc model selection testing the effects of fragment area on the structure of avian seed-dispersal networks. Area corresponded to the logarithmic scale of hectares per fragment. Intensity corresponds to the average number of interactions per species and was used as a fixed parameter to control for differences in sampling effort among studies. Only models with delta AICc < 2 were selected as plausible explanations for the observed patterns (see Table S1 for full model selection). *w*AICc gives an estimate of the probability of that given model to be the best choice under the AICc criteria. *r*^2^ gives an estimation of the variance explained by the optimal model while the coefficient *t* corresponds to the importance of that parameter within the model (*t* > 2 indicates the coefficient is significant with > 95% confidence). ‘+’ means the parameter was included in the model but had no influence in the final results.

| **Network metric** | **Area(log)** | **Intensity** | **Forest type** | **df** | **AICc** | **ΔAICc** | **w*i*** |
| --- | --- | --- | --- | --- | --- | --- | --- |
| N bird species | 0.08 | 0.17 |  | 4 | 25.5 | 0 | 0.75 |
|  |  | 0.21 |  | 3 | 27.7 | 2.19 | 0.25 |
|  |  | 0.15 | + | 6 | 36.0 | 10.47 | <0.01 |
|  | 0.08 | 0.13 | + | 7 | 40.9 | 15.37 | <0.01 |
| N plant species | 0.14 | -0.23 |  | 4 | 41.1 | 0 | 0.84 |
|  |  | -0.17 |  | 3 | 44.5 | 3.33 | 0.16 |
|  |  | -0.32 | + | 6 | 51.3 | 10.13 | <0.01 |
|  | 0.19 | -0.37 | + | 7 | 53.7 | 12.61 | <0.01 |
| N interactions | 0.12 | 0.70 |  | 4 | 26.7 | 0 | 0.96 |
|  |  | 0.75 |  | 3 | 33.5 | 6.75 | 0.33 |
|  |  | 0.65 | + | 6 | 36.9 | 10.21 | <0.01 |
|  | 0.11 | 0.63 | + | 7 | 40.1 | 13.38 | <0.01 |
| Number of links | 0.15 | 0.19 |  | 4 | 36.3 | 0 | 0.86 |
|  |  | 0.17 | + | 6 | -9.64 | 4.37 | 0.10 |
|  |  | 0.26 |  | 3 | -17.07 | 5.89 | 0.05 |
|  | 0.02 | 0.16 | + | 7 | -9.59 | 10.92 | <0.01 |
| Nestedness | -1.13 | 6.75 |  | 4 | 109.9 | 0 | 0.72 |
|  |  | 6.25 |  | 3 | 112.3 | 2.36 | 0.22 |
|  |  | 7.62 | + | 6 | 115.2 | 5.25 | 0.05 |
|  | -0.77 | 7.79 | + | 7 | 120.6 | 10.73 | <0.01 |
| H2’ specialization |  | 0.08 |  | 3 | 4.1 | 0 | 0.68 |
|  | -0.02 | 0.09 |  | 4 | 5.6 | 1.53 | 0.31 |
|  | 0.11 |  | + | 6 | 12.6 | 8.49 | 0.10 |
|  | 0.11 | 0.01 | + | 7 | 10.2 | 15.15 | <0.01 |

**Fig. S1.** Frequency distribution of the number of events per unique combination of plant-bird species performing seed-dispersal interactions in the tropical fragments of the Atlantic Forest.

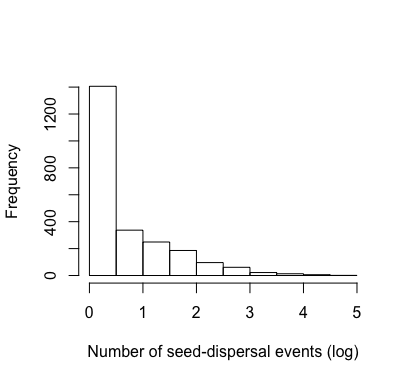
